# Phase Separation Clustering of Poly Ubiquitin Cargos on the Ternary Mixture Lipid Membranes by Synthetically Cross-Linked Ubiquitin Binder Peptides

**DOI:** 10.1101/2024.08.17.608403

**Authors:** Soojung Kim, Kamsy K. Okafor, Rina Tabuchi, Cedric Briones, Il-Hyung Lee

## Abstract

Ubiquitylation is involved in various physiological processes in our bodies such as signaling and vesicle trafficking, thus Ubiquitin (UB) is an important medical target system of interest. Polymeric addition of UB allows cargo molecules to be recognized specifically by multivalent binding interaction with UB binding proteins leading to various downstream processes. Recently, implication of protein condensate formation by Ubiquitylated proteins have been reported in many independent UB processes suggesting its potential role in governing the spatial organization of Ubiquitylated cargo proteins. We created modular polymeric UB binder motifs and polymeric UB cargos by synthetic bioconjugation and protein purification. Giant unilamellar vesicles with lipid raft composition were created to reconstitute polymeric UB cargo organization on the membranes. Fluorescence imaging was used to observe the outcome. We found that polymeric UB cargos clustered on the membranes by forming phase separation codomains in interaction with multivalent UB binder conjugate. This phase separation was valency dependent and was strongly correlated to their potency to form protein condensate droplets in solution. Multivalent UB binding interactions showed a general trend toward the formation of phase separated condensates, and the resulting condensates were either in liquid-like or solid-like state depending on the conditions and interactions used. It implies that polymeric UB cargos on the plasma and endosomal membranes may use the codomain phase separation to assist clustering of UB cargos on the membranes for cargo sorting. Our work also shows that model systems of such phase behavior can be created by modular synthetic approach that can potentially be used to further engineer biomimetic interactions in vitro.

**Graphical abstracts (TOC):** 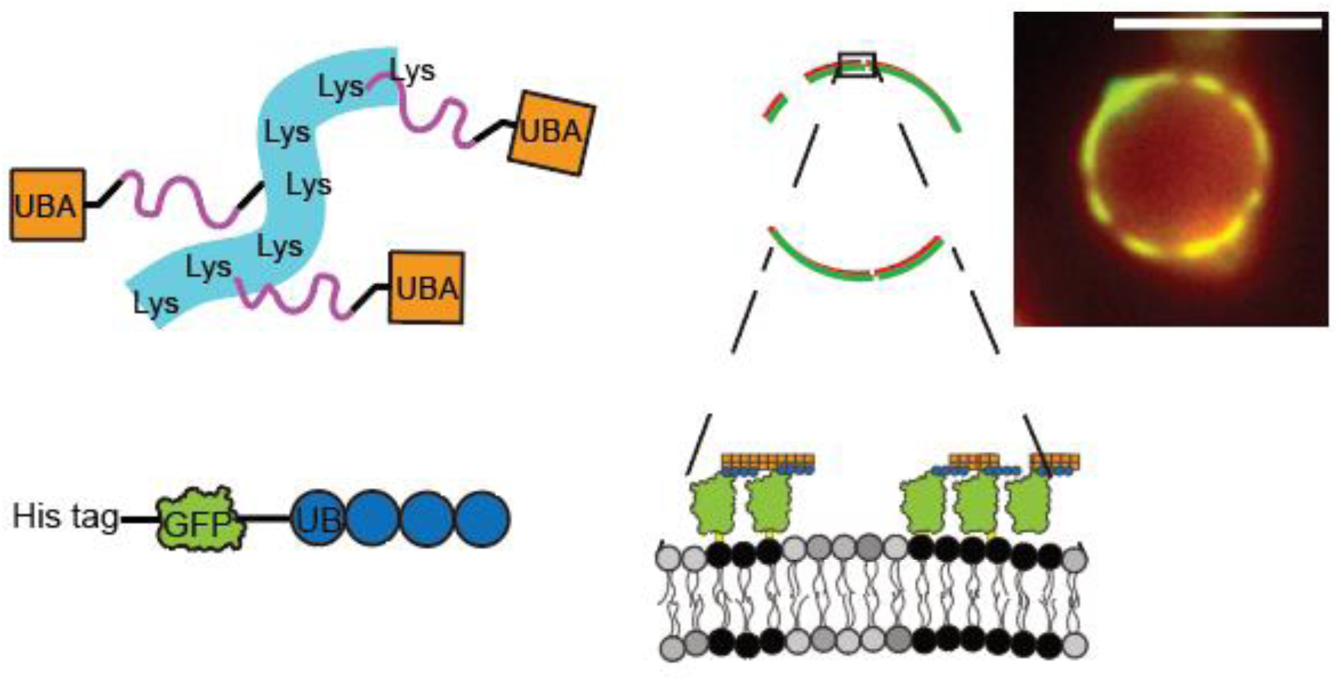

## 1. Introduction

Ubiquitylation is an enzymatic addition of a small protein, Ubiquitin (UB), to a target protein. As the name implies, Ubiquitylation is present ubiquitously in various physiological processes such as signaling^1^, DNA repair,^2^ and receptor trafficking.^3^ Due to its general and versatile usage in physiological processes of many organisms^4, 5^ including humans, the biochemistry of UB has been an intense area of research. For example, in endoplasmic reticulum associated protein degradation, (ERAD) misfolded proteins are Ubiquitylated and destined for degradation.^6, 7^ Maintaining such quality control systems is essential for cell survival. Cancerous cells are known to overburden the ERAD system, thus Ubiquitylation and its involvement in ERAD has been of great clinical interest as a therapeutic target to treat cancers.^8^ In vesicle trafficking of plasma membrane cargos, many trafficking proteins such as Clathrin^9^ and Endosomal complex required for transport (ESCRT)^10, 11^ family proteins are known to have UB binding domains to selectively bind to the UB bearing cargos for degradation or recycling of the target membrane proteins.

Recently, several groups have reported the relevance of liquid-liquid phase separation or condensate formation in UB involving processes, especially involving poly-Ubiquitin (polyUB), where UB addition is repeated in a polymeric manner allowing specific recognition by multivalent binding partners.^12, 13^ Multiple independent UB systems have been reported regarding their capability to form protein condensates. The reported systems include Rad23B in proteasomes,^14^ Ubiquilin in stress granules,^15, 16^ p62 in selective autophagy,^17^ ESCRT-0 in vesicle trafficking,^18^ and NF-κB essential modulator (NEMO) in NF-κB signaling.^19^ It suggests that condensate formation is an inherent property of the UB system and plays an important role in UB involving processes. Protein condensates can work as membranless organelles by enriching a subset of proteins inside the condensates.^20, 21^ Such enrichment may enhance the kinetic rate of biochemical reactions catalyzing the processes.^22, 23^ When such condensate formation happens on the lipid membranes involving membrane proteins, the condensate may cluster membrane proteins^24, 25^ and such clustering can enhance the kinetic rate of recruiting the downstream proteins involved in the processes.^26, 27^ Therefore, understanding the condensate formation involving UB is of great significance.

In this report, we used a synthetic modular approach to create polyUB and poly-UB binder proteins via a designer artificial protein construct and synthetic peptide crosslinking. We used wide field fluorescence and difference interference contrast (DIC) microscopy to observe the outcome of condensate formations from various combinations of the UB proteins. We show that membrane protein cargos, bearing the UB, may cluster via phase separation. Previous reports showed that a protein condensate formation and lipid raft formation may collaboratively form codomains on the membranes.^28–30^ We found polyUB cargos can form similar codomains when tested with synthetic giant unilamellar vesicles (GUV), mimicking the ternary compositions of the mammalian plasma membranes.^31^ We also show that such codomain formations are closely tied to the inherent property of polyUB proteins to form a phase separated protein condensate that may be in the final state of fluidic or solid phase. It suggests that such valency dependent cargo codomain formations may play an important role in the cargo clustering step of the vesicle trafficking, and a modular synthetic system can be used as a model system to study the physiological process systematically, and to design biomimetic systems in vitro.

## 2. Methods

### 2.1 GUV Preparation

All lipids used were purchased from Avanti Polar Lipids Inc., and stored in chloroform at -20°C. GUVs were created by the gentle hydration method.^32, 33^ The ternary mixture GUVs were comprised of a mixture of 1,2-Dioleoyl-sn-glycero-3-phosphocholine (DOPC), 1,2-Dipalmitoyl-sn-glycero-3-phosphocholine (DPPC) and Cholesterol. For homogenous GUVs, 35 mol% DOPC, 19.8 mol% DPPC and 35 mol% Cholesterol were used with added functional lipids of 0.2 mol% Texas Red-1,2-Dihexadecanoyl-sn-Glycero-3-Phosphoethanolamine (TR-DHPE, Invitrogen) and 10 mol% Nickel bound 1,2-Dioleoyl-sn-glycero-3-[(N-(5-amino-1-carboxypentyl)iminodiacetic acid)succinyl] (Ni-DOGS). Between 200– 500μg of lipid mixture was loaded into a round-bottom glass flask and insufflated with high purity nitrogen to generate a thin lipid film, which was subsequently desiccated in a vacuum chamber for at least 1 hour at room temperature, to remove residual chloroform. Lipid films were gently hydrated by addition of 1mL of 320 mM aqueous sucrose solution and incubation at 37 °C for 16–19 hours. GUVs were harvested post-incubation by centrifugation at 12,000*g* for 5 min to remove aggregation and was stored at 4°C to be used within a day.

### 2.2 GUV-Protein Sample Preparation for Experimentation and Imaging

Circular cover glass (Thickness #1, Fisher scientific) was immersed in a 1:1 mixture of clean water and isopropanol, then cleaned by bath sonication for 30 min followed by rinsing with ultrapure water. The coverglass was assembled into an Attofluor sample chamber (Invitrogen) with a O-ring to limit the total sample volume to 100-200 μL. The cover glass was surface passivated with 200 μl of 5 mg/ml bovine serum albumin (BSA) solution for 30 min. Residual BSA was rinsed and buffer exchanged into the Hepes buffer solution. (20 mM Hepes, 150 mM NaCl, pH 7.4) All solutions were created with clean water that was reverse osmosed, filtered, and ion exchanged multiple times that was finalized with the Milli-Q filtration unit. (Millipore sigma) 1-10 μL of GUV solution to ensure countable number of vesicles per images was added into the chamber and allowed to equilibrate for 10 min. Representative z-stack images were captured at multiple unique positions, to wholly characterize the GUVs present based on their shape, lamellarity and size. For anchoring of the UB-Green fluorescence protein (GFP) cargo to the membranes, His-tagged UB-GFP cargo proteins were added as final ∼4 µM concentration. Any step-wise addition was done by adding at least 10 vol% of the current liquid volume to ensure homogeneous mixing. After 30 min incubation, z-stack images were taken to monitor the successful binding of cargos. Lastly, UBD-Conjugate was added to cause interaction with UB cargos by adding ∼17 µM by UBD monomer concentration. (∼120 µM by Lys monomer concentration) 30 min was given during which time, time-lapse images were taken to monitor kinetic changes as needed. After the incubation, many z-stack images were sampled as final state images. All experiments were done at room temperature 22(±1) °C. Images were analysed for phase separation state of individual vesicles that are well defined as unilamellar vesicles.

### 2.3 Protein condensate formation

For solution phase condensate formation, UB-GFP cargos and UBD Conjugate were mixed in an Eppendorf tube at ∼30 µM for UB-GFP cargos and ∼200 µM for UBD Conjugate by the UBD monomer concentration (1.5 mM by Lys monomer concentration) to incubate for at least 10 min at room temperature. The mixture was imaged directly by adding a drop of liquid on a clean coverglass assembled into the sample chamber. Many images in fluorescence and DIC were collected as final state outcome. Other mixtures were incubated as 200 µM flexible UB6 + 100 µM NEMOUBAN6 for UB6/NEMOUBAN6 system, and 70µM UB6 + 120µM UBA6 for UB6/UBA6 system. For UB6/NEMOUBAN6 and UB6/UBA6, a fraction of UB proteins were labelled with an organic fluorescent dye Sulfo-Cy5 (Lumiprobe) for fluorescence signal.

### 2.4 Imaging Conditions

GUVs were imaged by a Nikon Ti2E-based inverted epifluorescence microscope system (Nikon, Japan) mounted on a vibration isolation table. The Nikon Apo 100x TIRF oil objective lens with numerical aperture of 1.49 was used with an sCMOS camera (Hamamatsu ORCA Flash 4.0, Hamamatsu, Japan) for image acquisition. An LED white light (Lumencor, Beaverton, OR), filtered by dichroic mirrors and optical filters to transmit selective wavelengths, was used to excite the GUV samples, emitting green or red fluorescence signals in the GFP or Texas Red channels, respectively. Micromanager was used for automatic acquisition of z-stack images from 1µm-20µm and the x, y positions were mechanically controlled to visualize unique z-sections of GUVs which were analyzed by FIJI (ImageJ) software. For DIC imaging, the same scope was used with contrast polarization filters and bright field light illumination in place.^34^

### 2.5 Protein Purification and Peptide Synthesis

The UBD Conjugate was synthesized by crosslinking.^35–37^ The synthesis was carried out in a 3-step reaction. Synthetic crosslinker bearing Malemide (Mal) group and N-Hydroxysuccinimide (NHS) ester in each end linked by 6 repetitions of poly ethylene glycol (PEG) was used. (BroadPharm, CA) Firstly, UBD peptide, produced to the purity of >99,9% (Genscript) was bound to the synthetic crosslinker using Cys-Mal reaction overnight by mixing 100µL of 1.0 mM peptide monomer with 100µL of 2.0 mM of the crosslinker (2x molar ratio) under the pH 7.4 20mM Hepes, 150mM NaCl, 2.5mM tris(2-carboxyethyl)phosphine (TCEP) condition at 4°C for 16–19 hours. Then the successfully crosslinked UBA peptides were bound to the PLL backbone by NHS ester-Lysine reaction overnight by adding with 25 µL of 4mg/mL Poly-L-Lysine (PLL, 4-20kD, MP Biomedicals) solution to the 0.44 mg/mL final concentration to incubate at 4°C for 16–19 hours. Finally, 5-10mM final concentration of 2-mercaptoethanol was added into the reaction to quench the reactive functional groups and stored at 4°C until used.

The sequence of the ESCRT-0 UBD peptide used was following, originally UB interacting motif from STAM1A of the ESCRT-0 complex.^38, 39^

GCSKEEEDLA KAIELSLKEQ RQQGGSWC

Tryptophan and cysteine were added for spectroscopy characterization and efficient crosslinking.

For purification of modularly designed proteins, recombinant plasmids (Genscript) were transformed into competent *E. Coli* for overexpression.^30, 32, 40, 41^ Used strains were BL21AI (Invotrgen) or BL21(DE3) where overexpression could induced with 0.5 mM IPTG and Arabinose (only for BL21AI). Flexible 6UB and 6UBA were expressed with BL21(DE3) and other proteins were expressed in BL21AI. *E.Coli* was grown and 37°C and overexpressed overnight at a lower temperature of 18°C. *E. Coli* cells were harvested by centrifugation and the retrieved pellets were resuspended in the Hepes buffer solution (20 mM Hepes, 150 mM NaCl, pH 7.4) and disrupted by high-pressure French press (Glen Mills, Clifton, NJ). Proteins were separated from *E. Coli* debris by centrifugation and purified through Ni-NTA affinity chromatography in a gravity column. Further purification was achieved by automated column chromatography (ÄKTA explorer, GH Healthcare) typically using a size exclusion chromatography of HiLoad Superdex75 or Superdex200 column. The purified proteins were characterized by SDS-PAGE and spectroscopy to store at -20°C. Protein sequence information can be found in the Appendix 2 of the Supporting Information.

## 3. Results and discussion

### 3.1. Purification and Synthesis of modular Ubiquitin proteins

To test the potency of the membrane UB cargos to cluster by phase separation, synthetic multivalent UB binder conjugate was designed and created. (Figure 1A) Briefly, UBD domain taken from the ESCRT-0 sequence, one of the UB binding motifs in the protein complex,^39^ was synthesized as a monomer peptide bearing Cys residues for Cys-Mal reaction. A synthetic crosslinker bearing Maleimide and NHS ester on each end was attached to the peptide first. This peptide-crosslinker was lastly attached to the PLL by Lys-NHS ester reaction. Multiple UBD peptides could bind to each PLL as every Lys was open for the crosslinking reaction, thus the reaction was limited by the number of reactable peptide-crosslinkers. Based on the series of characterization by size exclusion chromatography and UV absorption spectroscopy, it was estimated that typical multivalent conjugate that participated in the reaction bears median 4 UBD peptides per conjugate spanning the molecular weight of 10-40kD when synthesized with the condition we used. (Appendix 1 of the Supporting Information) By using this bioconjugate approach, we can test the multivalent interaction of interest at its purest form as a model system that can be potentially modified for further studies. This way, we are not limited by the purification with the mammalian cell expression^42^ of every component involved in the interaction which is often required to study a specific UB interaction system, and this may serve as a stepping stone toward the synthesis of biomimetic system utilizing the polyUB interaction in vitro. Various polyUB constructs were also designed as model membrane cargos with Ubiquitylation. (Figure 1B) GFP served the model cargo proteins while providing necessary fluorescence signal for microscopy. They were *E.Coli* expressed and purified by a series of chromatography. His-tag was used for affinity purification and was preserved for membrane binding by the his-Ni chelation with Ni-DGS lipids.

**Figure 1.**
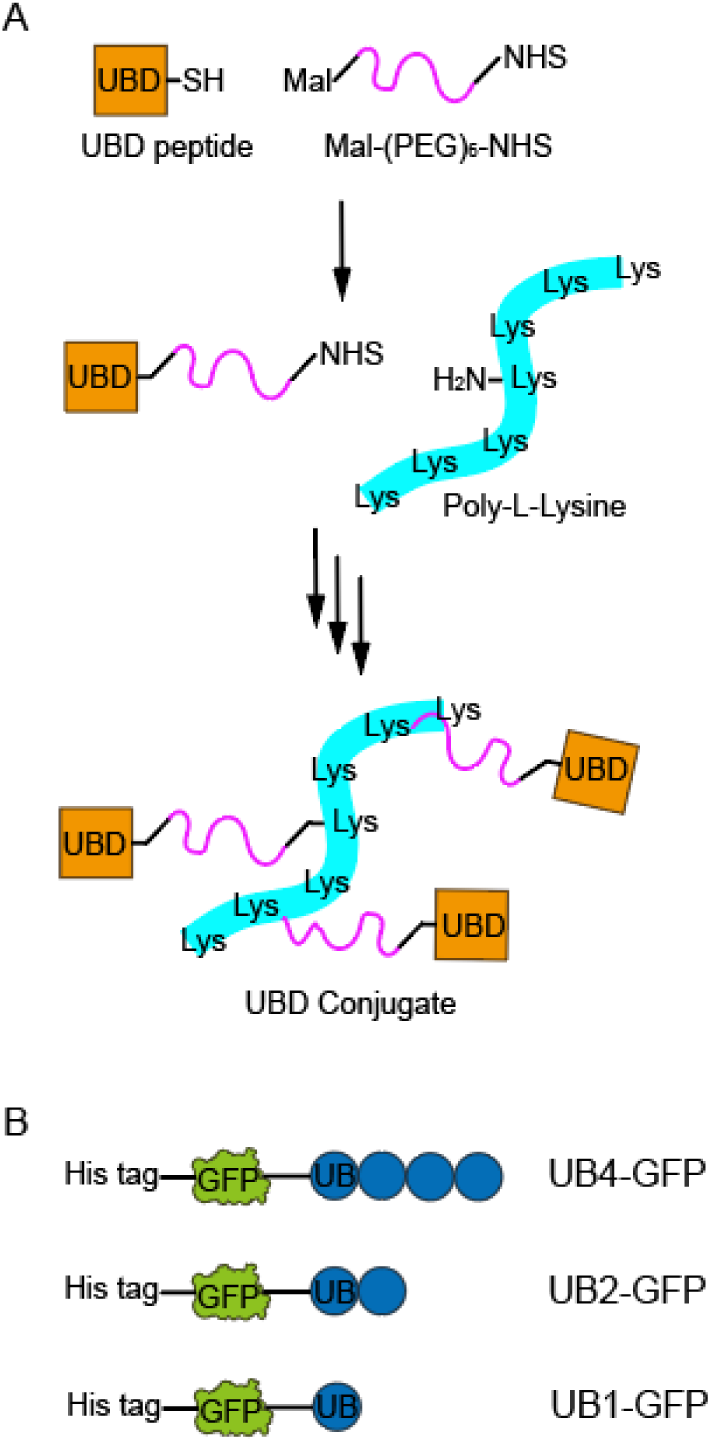
Modular UB proteins. (A) Synthetic UBD conjugate was created by crosslinking multiple UBD peptides into a PLL by the synthetic crosslinkers. (B) polyUB membrane cargos were created by purifying GFP bearing linear polyUB with varying lengths.

### 3.2. polyUB membrane protein cargos phase separate into lipid-protein codomains on the ternary mixture GUVs

To study the implication of phase separation in polyUB membrane cargo clustering using the modular synthetic proteins, we reconstituted the UB4-GFP cargo on the lipid membranes mimicking the composition of the physiological plasma membranes where saturated chain lipids, unsaturated chain lipids, and cholesterol coexist.^31, 43^ GUVs with the ternary mixture composition including DOPC, DPPC, and cholesterol were created. (DOPC 35.0 mol%, DPPC 19.8 mol%, Cholesterol 35.0 mol%, Ni-DGS 10.0 mol%, TR-DHPE 0.2 mol%) 10 mol% of what is supposed to be DOPC, when only the ratio of ternary mixture is considered, was replaced into a functional head group lipid Ni-DGS which was used to permanently anchor the UB4-GFP cargo to the lipid membranes. (see Supporting Information Figure S2 for negative control data.) 0.2 mol% TR-DHPE was introduced as a fluorescent reporter for the lipids. The GUVs were first introduced to the sample chamber to observe by fluorescence imaging, then the UB4-GFP cargos were bound to the membranes, and finally UBD conjugate was introduced to cause binding interaction between UB4-GFP and UBD Conjugate. (Figure 2A) Each step was examined via multi-color fluorescence imaging to sample many cases of vesicles. The resulting phase states of vesicles were statistically analyzed for each stage of incubation.

**Figure 2.**
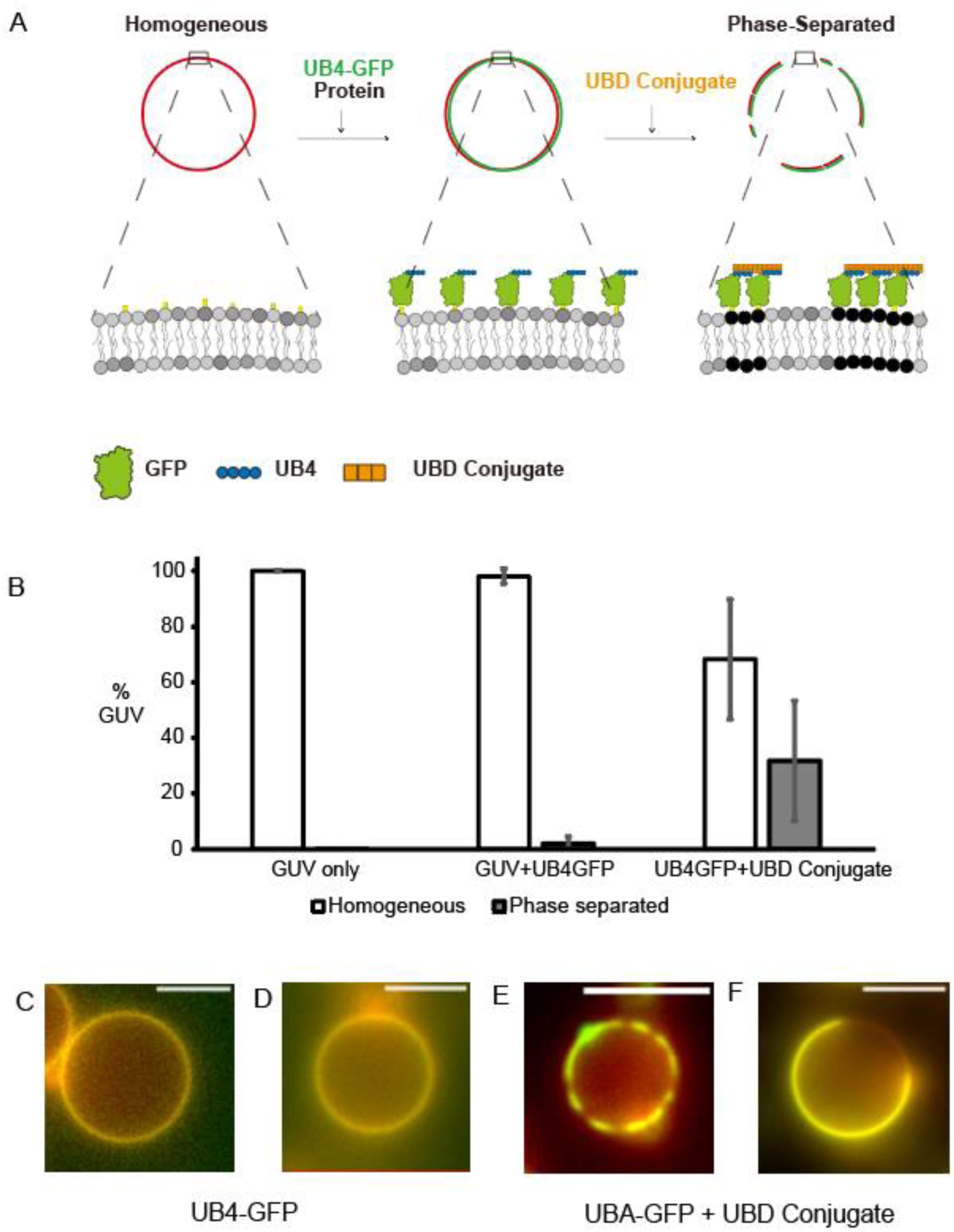
Phase separation of polyUB cargo on the membranes by polyUB-UBD Conjugate interaction. (A) Schematic of the lipid–protein interaction transitioning from homogeneous to phase-separated after introducing UB4-GFP and UBD Conjugate. The lipid composition was DOPC 35.0%, DPPC 19.8%, cholesterol 35%, Ni-DGS 10%, and TR-DHPE 0.2%, and incubation conditions of the proteins were 1 μM SUMO3-GFP and 1 μM SUMO10 and SIM10 each. (B) Statistical distribution of the resulting phase behavior of GUVs. Error bars represent standard deviations from each image taken. Statistical analysis from 25 image stacks taken from 3 independent experiments. (C, D) Example images of GFP cargo fluorescence (green) overlapped with Texas Red lipid fluorescence (red), maintaining homogeneous behavior after introducing UB4-GFP protein. (E, F) Example images showing phase-separated behavior after introducing UBD Conjugate. Scale bars are 5 μm.

Ternary mixture lipids are known to separate into liquid ordered (lo) and liquid disordered (ld) binary lipid domains at specific conditions.^31, 44–47^ This composition GUVs we used mostly remained homogeneous at room temperature, and this distribution did not change after UB4-GFP cargo was introduced on the membranes. However, once UBD Conjugate was introduced to interact, it caused about 30% of the vesicles, on average, to form phase separated cargo domains. (Figure 2B) Example images of homogeneously distributed and phase separated cargos are also shown in Figure 2C-F. As shown in the examples, the cases of UB4-GFP cargos clustering into cargo enriched space leave the rest of the cargo depleted space readily visible only after introducing the UBD Conjugate. This is because multivalent interaction between UB4-GFP and UBD Conjugate caused cargos to phase separate forming liquid condensate on the membranes.^48^

Many previous studies suggested that the driving force of the ternary mixture lipid membranes to form lo and ld domains and the driving force of the proteins on the membranes to form protein condensate domains can collaboratively cause the formation of codomains where protein domains and lipid domains align in space.^28–30, 32^ Our result suggests that this principle applies to the polyUB cargos on plasma membrane mimicking ternary mixture lipid membranes as well. This is important because it implies that polyUB cargos on the membranes are poised at a condition where they can cluster into phase domain easily to aid the downstream processes such as vesicle trafficking. It potentially implies that plasma membrane and endosomal cargo sorting step is assisted by the compositions of the membranes to better organize the cargo proteins on the membranes in space.

In our previous study using SUMO cargo proteins, we argued that when the collaborative codomain formation is in play, the phase separation tends to promote domain formation while there exists the opposing force of steric pressure of having high density cargos on the membranes.^32, 49^ These two opposing forces are at a “tug of war” to determine the final statistical distribution of cargo domain separation. In the case of SUMO cargo proteins, the high density cargo condition caused by the 10 mol% Ni-DGS functionalization lipids incubated with comparable concentrations of SUMO3-GFP caused the resulting cargo distribution to be exclusively homogeneous while lower Ni-DGS concentrations of 5, 1 mol% reduced the steric pressure to increase the distribution of phase separation.^32^ We found it interesting that this UB4-GFP + UBD Conjugate system at a highly dense condition, as 10 mol% Ni-DGS, ended up resulting in a significant amount of phase separated cargos. We were able to find some cases of kinetic changes that imply the existence of aforementioned “tug of war” between domain formation and reversing steric pressure (Supporting Information Figure S3), but the effect of steric pressure was much less pronounced for polyUB – UBD Conjugate interaction. Steric pressure of structured proteins is dependent on the size of the proteins interacting,^49^ so we speculate that this discrepancy between the UB and SUMO systems may be due to the difference is sizes of the binder protein used where UB Conjugate was 20-40kD and SUMO10 and SIM10 used in SUMO study were 119 kD and 34 kD each.

### 3.3. Codomain formation is valency dependent

After observing the potency of polyUB cargos on the membranes to form codomain by its interaction with multivalent binder UBD conjugate, we sought to systematically study the valency dependence of the phase separation. It is widely recognized that often favorable multivalent interaction is the key to promote protein condensate by shifting the equilibrium toward the formation of a dense condensate structures.^40, 41^ In physiological processes, Ubiquitylation happens at various lengths and morphologies,^50, 51^ so the dependence on valency is important.

Firstly, a negative control experiment was performed with a monomer UBD binder. Identical concentrations of monomer UBD binder peptides were introduced in the last step of the experiment instead of the UBD conjugate. The identical GUV composition and UB4-GFP concentration were used as shown in the original experiment of Figure 2 other than the use of the UBD monomer. In contrast to the original result with UBD Conjugate (Figure 3A), UBD monomer resulted in practically no change after its introduction to the system. (Figure 3B) There are identical number of UBD motifs in solution, thus it is the multivalency in binding interaction that caused the difference between codomain formation and no significant change at all.

**Figure 3.**
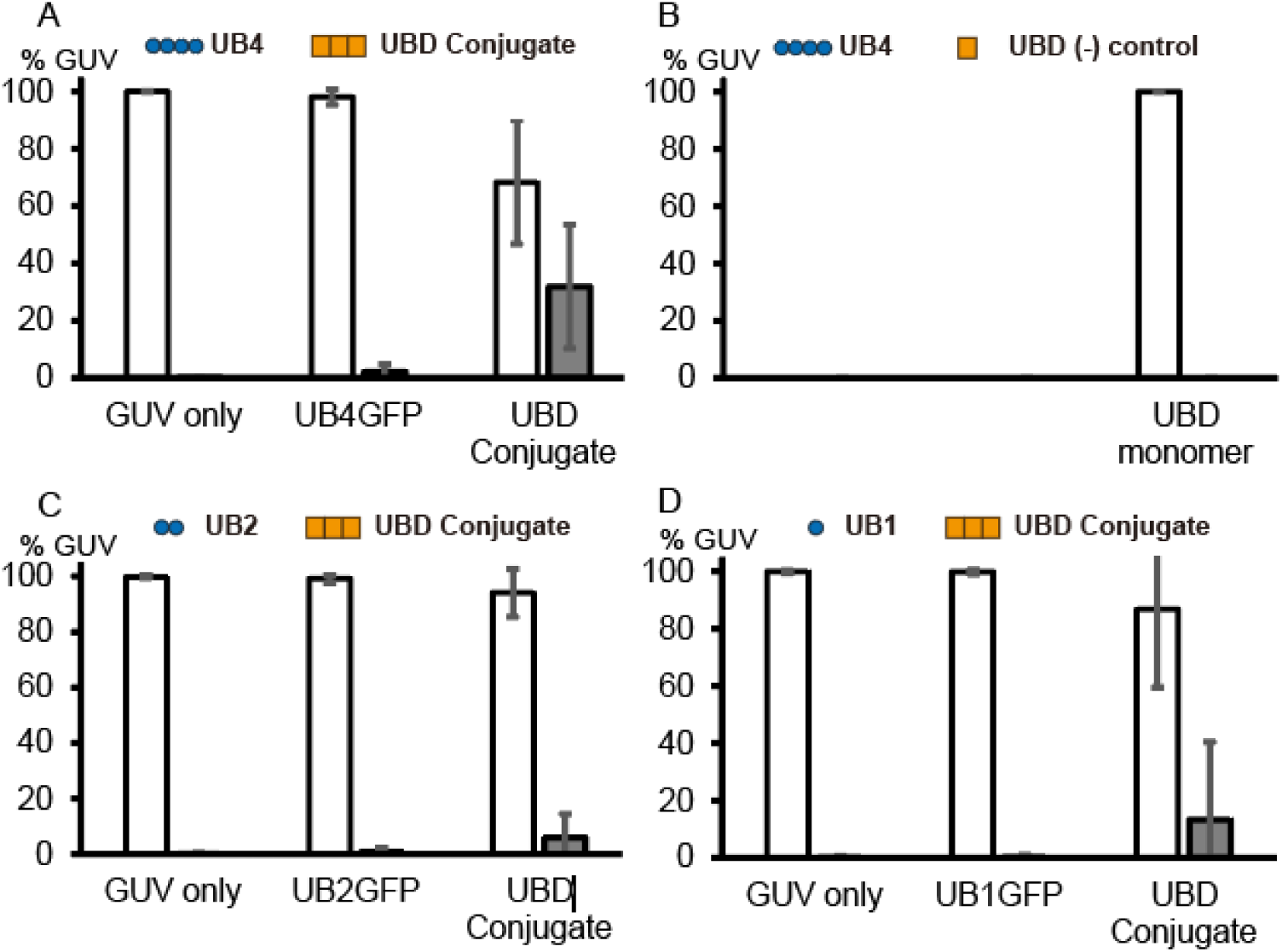
Statistical analysis of phase separation state of the cargo vesicles. (A) UB4-GFP and UBD Conjugate experiment. Reproduction Figure 2B. (B) UB4-GFP and UBD monomer experiment. Identical to the experiment A except that UBD monomer was used as negative control of the binding multivalency. GUV only and UB4GFP omitted to avoid redundancy with A. (C) UB2-GFP and UBD Conjugate experiment. (D) UB1-GFP and UBD Conjugate experiment. Error bars represent standard deviations from each image taken. Statistical analysis were from at least 25 image stacks taken from 3 independent experiments for each.

We then performed similar series of experiments by varying the length of Ubiquitylation of the membrane cargos. UBD Conjugate was kept constant as a multivalent binder while varying the length of polyUB by 2 and 1. As shown in Figure 3CD, shorter UB cargos had much less tendency toward phase separation as evidenced by ∼10% or less in final statistics of phase separated cargo vesicles. It matches with general expectations where greater multivalency promotes the formation of phase separated protein condensates, and polymeric Ubiquitylation has a distinct potency to form such codomain on the membranes compared to mono- or di-Ubiquitylation of cargos. It may suggest a way to affect the fate of membrane protein cargos poly Ubiquitylation.

### 3.4. Dynamic changes in time lapse images confirm that the clustering domain formation is collaborative between lipids and proteins

To obtain the kinetic trace during the statistical change of the phase separation state, we performed time lapse imaging analysis for the UB4-GFP and UBD Conjugate interaction. Multi-color images were taken every 2 min after the initial introduction of the UBD Conjugate to the sample. Time zero was defined as the very time UBD Conjugate was introduced. Figure 4 shows two time-lapse examples of the typical change from homogeneous to phase separated state after the incubation started. As is evident in the GFP channel images, that represent the spatial distribution of the UB4-GFP cargos on the membranes, initially homogeneous fluorescence distribution start developing bright and dark spots across the membranes that indicate binary separation into the phase domains. The separation domain may show the behavior of fluctuation and coarsening in time scale of min as they are fluidic domains but eventually maintains the state of separated state. When TR channel images are compared, which represents the behavior of the lipids,^52^ it is evident that lipids separation spatially overlap with protein cargo separation as bright/dark regions in TR channels reproduces bright/dark regions in GFP channels precisely. It is worth noting that lipids and protein cargos may move independently in a hypothetical scenario of proteins and lipids not interacting strongly. It is the formation of codomains between lipids and proteins by a favorable intermolecular interaction that force them to move correlated in space.

**Figure 4.**
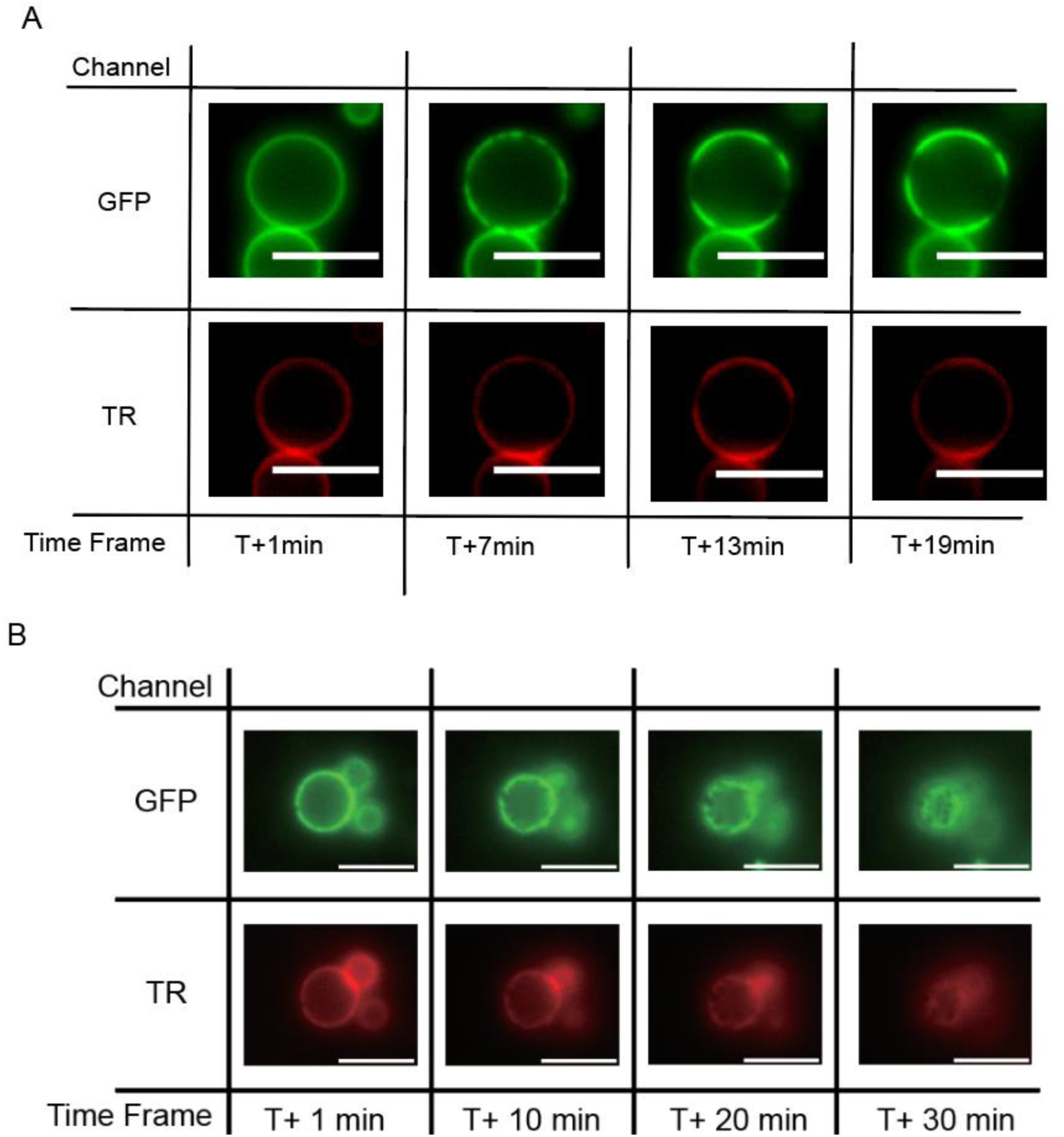
Time lapse images show dynamic changes of the phase separation states. (A) An example time lapse taken at T+1min, 7min, 13min, and 19min. Time zero is defined as a time the UBD Conjugate was added to the sample with UB4-GFP cargo on ternary mixture GUVs. What is originally homogeneous becomes intermittently clustered in T+7min, then coarse into phase separated domains in T+13min. TR signal shows lipids follow the localization of the protein cargos. (B) An example time lapse taken at T+1min, 10min, 20min, and 30min. Similarly, originally homogeneous vesicle develop dark regions as a result of phase separation. Focus moved up a bit toward the later time point better visualizing phase separation of the upper top portion of the vesicle. Lipids and protien signals in two channels generally agree in space. Scale bars are 5 µm.

### 3.5. polyUB proteins have potency of forming protein condensates in solution at various states and the potency is strongly correlated to the codomain formation on the membranes

Multiple recent literatures reported the potency of UB proteins to form protein condensate in solution.^14–16, 18, 19, 53^ We tested the protein condensate formation in solution without the membranes at higher concentrations than the membrane experiments (>30µM each). UB cargo proteins and UBD Conjugate were mixed in solution and allowed enough time to interact for at least 10 min at room temperature and the outcome was observed by microscopy. As shown in Figure 5A, UB4-GFP and UBA Conjugate readily formed many condensates with round droplet shapes greater than 1 µm in diameter. The droplets were enriched with UB4-GFP as evidenced by the bright fluorescence signal, and the droplets were clearly visible by DIC imaging. When identical incubation was attempted with the same total concentrations of UBD monomers that lack multivalency, there was no visible entities detectable in solution indicating the absence of condensate forming interactions. UB2-GFP and UB1-GFP, when tested for the same interactions, were able to form some visible droplets albeit smaller and less in number. Most prominently, the enrichment of UB-GFP cargos inside the droplets were several times less compared to the UB4-GFP as estimated by the fluorescence intensity (Supporting Information, Figure S4). It suggests that UB4-GFP, with its greatest multivalency, has the strongest potency to form protein condensate and to enrich it with the most cargo molecules that correlate to the statistical result of forming most phase separated cargo vesicles on the membranes in Figure 3.

**Figure 5.**
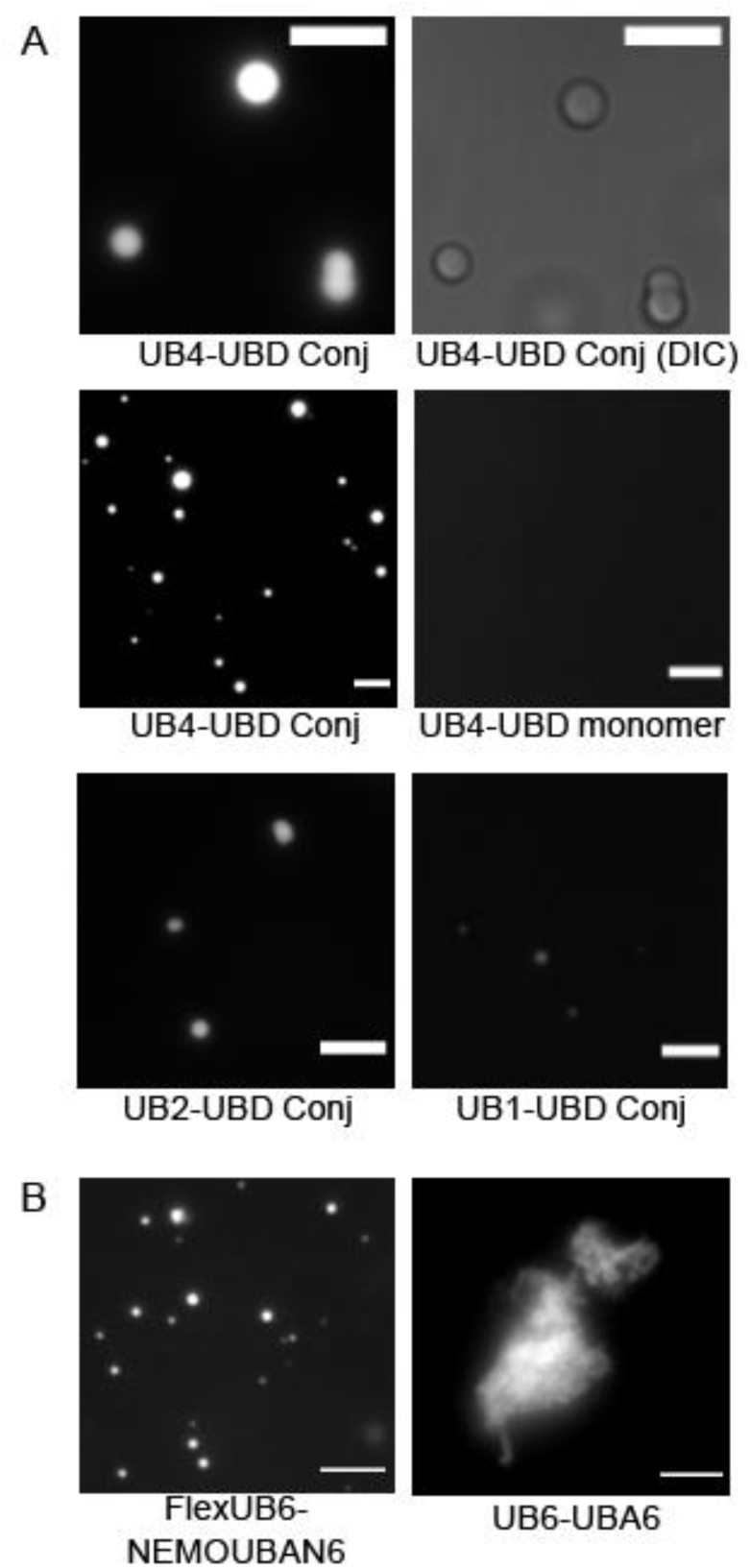
Protein condensate formation of UB proteins in solution. (A) Example condensate images between UB-GFP in varying lengths (UB4, UB2, and UB1) and UBD conjugate, UBD monomer in fluorescence and DIC. The fluorescence and DIC images in the first raw are matching images. (B) Example condensate images formed between other polyUB and UB binder proteins. FlexUB6 and NEMOUBA6 form liquid-like droplets while UB6 and UBA6 form solid-like structures. Scale bars are 5 µm.

We also tested numerous other UB and UB binder proteins regarding their potency to form protein condensate in solution. Figure 5B shows example images of solution phase behavior of other UB proteins. (see Supporting Information Figure S5 for schematics of the proteins.) Flexible-UB6, labeled by an organic fluorescence dye which is a construct of polyUB linked by flexible GGS spacer between UBs, when interacted with another modular protein of NEMOUBAN6, a six times repetition of UBAN domain from NEMO proteins, showed clear formation of round condensate droplets comparable to the one formed between UB4-GFP and UBD conjugate. This system used a different UB binding domain and purification of modular construct instead of peptide synthesis. It shows that the modular repetition of UB and UB binding multivalency can reproduce such condensate formation behavior regardless of the exact nature and structure of the multivalent binding interactions.

It is very important to note that not all modular interactions involving multivalent interaction between polyUB and UB binder proteins would lead to liquid-like phase separation.^54, 55^ PolyUB proteins may form condensate at solid-like state as shown for the case between UB6 (linear polyUB with 6 times UB repetition labeled with an organic dye) and UBA6 (linear 6 times UBA domain^56^ repetition purified by overexpression in E.Coli). Solid-like structures tend to have irregular shapes instead of round shapes, and they tend to grow into a gigantic rock-like spiky structures greater than 10 µm in length. (Figure 5B) It is well known in the field of phase separation of protein condensate that at a certain subregion of phase space, the proteins may form solid-like structures.^54, 55, 57–59^ Its relevance in physiological interaction of polyUB cargos is to be investigated, but it is important that they have potency to form such solid-like structure at certain conditions of interaction.

## 4. Conclusion

In this study, we report that polyUB cargos on the ternary mixture lipid membranes that mimic the composition of the plasma membranes can collaboratively cluster into phase separated domains that are dependent on the multivalent interaction with UB binding proteins. We constructed a simple model system to test this by modularly designed synthetic UBD Conjugate and polyUB cargos at varying lengths. The membrane clustering is closely correlated to their potency to form protein condensate in solution. polyUB proteins in multivalent interaction has a general potency to form condensate, but the resulting state of the condensate could be liquid-like or solid-like depending on the conditions and the proteins involved.

PLL is a commonly available peptide these days so the strategy of using PLL as a backbone to attach multiple binding motifs opens a possibility to create and engineer various similar synthetic systems^37, 60, 61^ to test multivalent binding interactions. The model can test interactions independent from the effect of the rest of the protein structures and is easily expandable to test fundamental intermolecular interactions that are crucial for the desired outcome. It can be applied to design in vitro biomimetic systems utilizing the condensate-forming interactions.^62, 63^ Caution should be taken though, that synthetic approach is missing potential allosteric effects of the physiological proteins that may be important to finely regulate each specific physiological interaction.^39, 64^ As we saw in this investigation, our UBD Conjugate was relatively small in size, potentially altering the opposing effect of steric pressure of the physiological proteins, thus the result should be interpreted within the limitation of the model system.

Even considering the limitation of the model system approach, it is very evident that polyUB cargos on the membranes have the potency to cluster by forming phase separated codomain assisted by the raft forming lipid compositions. It suggests that poly Ubiquitylation may be used to promote spatial sorting of cargos on the plasma and endosomal membranes in vesicle trafficking processes. Multivalent binding and poly Ubiquitilation are very common in UB systems, thus their synergetic interaction with lipid membranes deserves attention in the related processes. ESCRT-0 protein complex, where the UBD domain was taken from, has multiple UB binding sites that were shown to form condensate in Yeast vacuoles,^18^ and many downstream ESCRT family proteins also possess UB binding motifs while capable of forming polymeric structures on the membranes for the purpose of remodeling the membranes.^10, 65^

Considering the increasing number of reports of case studies on polyUB based condensates, it is evident that such condensate formation plays a role in physiological processes. We learned that UB and UB binder proteins form not only liquid-like condensate but also solid-like structure in a reproducible manner. There have been no systematic studies regarding what leads to formation of solid-like structure instead of fluidic condensate, thus systematic investigation of the phase diagram of UB proteins is a potential future subject of study. Our working hypothesis is that highly dense multivalent interact in space beyond certain binding affinity causing solid-like structure formation by “a bit too tight” packing of the proteins that need further scrutiny in future projects.

## Abbreviations

DIC: Differential interference contrast microscopy
DOPC: 1,2-Dioleoyl-sn-glycero-3-phosphocholine
DPPC: 1,2-Dipalmitoyl-sn-glycero-3-phosphocholine
ESCRT: Endosomal sorting complex required for transport
GFP: Green fluorescence protein
GUV: Giant unilamellar vesicle
Mal: Maleimide
NEMO: NF-kappa-B essential modulator
NHS: N-Hydroxysuccinimide
Ni-DGS: Nickel bound 1,2-Dioleoyl-sn-glycero-3-[(N-(5-amino-1-carboxypentyl)iminodiacetic acid)succinyl]
PEG: Poly ethylene glycol
PLL: Poly-L-Lysine
polyUB: Poly Ubiquitin
TCEP: tris(2-carboxyethyl)phosphine
TR-DHPE: Texas Red-1,2-Dihexadecanoyl-sn-Glycero-3-Phosphoethanolamine
UB: Ubiquitin
UBA: Ubiquitin association domain
UBAN: Ubiquitin-binding domain in ABINs and NEMO
UBD: Ubiquitin binding domain

## Author contributions

Conceptualization and design of the project, I.-H.L.; Experimental work and data collection, S.K., K.K.O., R.T., C.B.; Data analysis, S.K., K.K.O.; Writing the original manuscript, I.-H.L., S.K., K.K.O.; Reviewing and editing the manuscript, S.K., K.K.O., R.T., C.B., and I.-H.L.

## Acknowledgements

The authors would like to thank Dr. Laying Wu of the Microscopy and microanalysis research laboratory at Montclair state university for the general microscope training and facility maintenance.

## Supporting Information

Supplementary materials including Figures S1-S5 and Appendix 1, 2.

