## Supporting Information for "Phase Separation Clustering of Poly Ubiquitin Cargos on the Ternary Mixture Lipid Membranes by Synthetically Cross-Linked Ubiquitin Binder Peptides"

and Il-Hyung Lee[1]*

Affiliations: [1]Department of Chemistry and Biochemistry, Montclair State University, Montclair, NJ 07043, USA, [2]Department of Biology, Montclair State University, Montclair, NJ 07043, USA

**The file includes:**

Appendix 1. Characterization of the synthetic UBD Conjugate

Appendix 2. Full sequence of the proteins used

S1-S5. Supplementary Figures

### Appendix 1. Characterization of the synthetic UBD Conjugate

We performed a series of experiments to characterize the synthesized UBD Conjugate. Since our synthesis relies on stochastic events of UBD peptides being crosslinked to the PLL backbone, the resulting UBD Conjugate is a mixture of n-mers that can be characterized by the average number of UBD peptides bound per backbone. Figure S1 shows the result of the characterization experiment including UV absorption spectroscopy (Nanodrop, ThermoFisher Scientific), size exclusion chromatography.

Firstly, we started by estimating the total amount of UBD peptide that successfully ended up binding to the backbone with molecular weight greater than 10kD. We introduced a tryptophan to the peptide sequence for UV absorption, thus UBD peptides selectively absorbed 280nm UV light while PLL and synthetic crosslinkers were invisible to UV absorption, which could be used as an advantage to selectively quantify the amount of UBADpeptides. We compared the 280nm absorption of the UBD Conjugate as it is, and also after centrifugation with 10kD threshold Amikon tube (EMD Millipore) to selectively retain UB Conjugate that is >10kD after 20 fold dilution. 280nm absorption was reduced to 10% of the original absorption before the filtration. This is not due to the change of concentration as the sample volume was kept constant. It suggests that about 90% of the UBA peptides fail to participate in the crosslinking reaction to make final molecular weight greater than 10kD. From this, we can reasonably assume that the majority of peptides do not end up being crosslinked to the backbone and they mostly remain as monomers. Thus the contribution of two cysteine residues causing crosslinking of multiple PLL backbone, although is a technical possibility, should be minimal.

Since even two UBD peptides being crosslinked to a 5kD PLL will have a molecular weight greater than 10kD, it can be assumed that any UBD Conjugate with meaningful multivalency that participated in the phase separation reaction should have molecular weight greater than 10kD. To learn the size distribution of the final UBD Conjugates, the sample was run by the Superdex S75 column by automated chromatography system using neutral hepes buffer. When compared to the standard calibration run by known molecular weight standards, it was estimated that the resulting UBD Conjugate has molecular weight distribution of 10-40kD where 25kD would be the median molecular weight.

The PLL we used for synthesis originally had molecule weight distribution of 4-20kD based on the manufacturer's characterization. If we assume 10kD of PLL backbone molecular weight, 10-40kD implies 0-8 mers per backbone with 4-mer being average for 25kD. (molecular weight of a peptide + synthetic crosslinker is 3.654kD)

### Appendix 2. Full sequence of the proteins used

*UBD peptide*

GCSKEEEDLA KAIELSLKEQ RQQGGSWC

*UB4-GFP*

MGMQIFVKTL TGKTITLEVE PSDTIENVKA KIQDKEGIPP DQQRLIFAGK QLEDGRTLSD

YNIQKESTLH LVLRLRGGMQ IFVKTLTGKT ITLEVEPSDT IENVKAKIQD KEGIPPDQQR

LIFAGKQLED GRTLSDYNIQ KESTLHLVLR LRGGMQIFVK TLTGKTITLE VEPSDTIENV

KAKIQDKEGI PPDQQRLIFA GKQLEDGRTL SDYNIQKEST LHLVLRLRGG MQIFVKTLTG

KTITLEVEPS DTIENVKAKI QDKEGIPPDQ QRLIFAGKQL EDGRTLSDYN IQKESTLHLV

LRLRGGCLEV LFQGPMVSKG EELFTGVVPI LVELDGDVNG HKFSVSGEGE GDATYGKLTL

KFICTTGKLP VPWPTLVTTL TYGVQCFSRY PDHMKQHDFF KSAMPEGYVQ ERTIFFKDDG

NYKTRAEVKF EGDTLVNRIE LKGIDFKEDG NILGHKLEYN YNSHNVYIMA DKQKNGIKVN

FKIRHNIEDG SVQLADHYQQ NTPIGDGPVL LPDNHYLSTQ SKLSKDTNEK RDHMVLLEFV

TAAGITLGMD ELYKENLYFQ GLEHHHHHH

*UB2-GFP*

MGMQIFVKTL TGKTITLEVE PSDTIENVKA KIQDKEGIPP DQQRLIFAGK QLEDGRTLSD

YNIQKESTLH LVLRLRGGMQ IFVKTLTGKT ITLEVEPSDT IENVKAKIQD KEGIPPDQQR

LIFAGKQLED GRTLSDYNIQ KESTLHLVLR LRGGCLEVLF QGPMVSKGEE LFTGVVPILV

ELDGDVNGHK FSVSGEGEGD ATYGKLTLKF ICTTGKLPVP WPTLVTTLTY GVQCFSRYPD

HMKQHDFFKS AMPEGYVQER TIFFKDDGNY KTRAEVKFEG DTLVNRIELK GIDFKEDGNI

LGHKLEYNYN SHNVYIMADK QKNGIKVNFK IRHNIEDGSV QLADHYQQNT PIGDGPVLLP

DNHYLSTQSK LSKDTNEKRD HMVLLEFVTA AGITLGMDEL YKENLYFQGL EHHHHHH

*UB-GFP*

MQIFVKTLTG KTITLEVEPS DTIENVKAKI QDKEGIPPDQ QRLIFAGKQL EDGRTLSDYN

IQKESTLHLV LRLRGGCLEV LFQGPMVSKG EELFTGVVPI LVELDGDVNG HKFSVSGEGE

GDATYGKLTL KFICTTGKLP VPWPTLVTTL TYGVQCFSRY PDHMKQHDFF KSAMPEGYVQ

ERTIFFKDDG NYKTRAEVKF EGDTLVNRIE LKGIDFKEDG NILGHKLEYN YNSHNVYIMA

DKQKNGIKVN FKIRHNIEDG SVQLADHYQQ NTPIGDGPVL LPDNHYLSTQ SKLSKDTNEK
RDHMVLLEFV TAAGITLGMD ELYKENLYFQ GHHHHHHHH

*UB6*

MQIFVKTLTG KTITLEVEPS DTIENVKAKI QDKEGIPPDQ QRLIFAGKQL EDGRTLSDYN
IQKESTLHLV LRLRGGMQIF VKTLTGKTIT LEVEPSDTIE NVKAKIQDKE GIPPDQQRLI
FAGKQLEDGR TLSDYNIQKE STLHLVLRLR GGMQIFVKTL TGKTITLEVE PSDTIENVKA
KIQDKEGIPP DQQRLIFAGK QLEDGRTLSD YNIQKESTLH LVLRLRGGMQ IFVKTLTGKT
ITLEVEPSDT IENVKAKIQD KEGIPPDQQR LIFAGKQLED GRTLSDYNIQ KESTLHLVLR
LRGGMQIFVK TLTGKTITLE VEPSDTIENV KAKIQDKEGI PPDQQRLIFA GKQLEDGRTL
SDYNIQKEST LHLVLRLRGG MQIFVKTLTG KTITLEVEPS DTIENVKAKI QDKEGIPPDQ
QRLIFAGKQL EDGRTLSDYN IQKESTLHLV LRLRGGMQIF VKTLTGKTIT LEVEPSDTIE
NVKAKIQDKE GIPPDQQRLI FAGKQLEDGR TLSDYNIQKE STLHLVLRLR GGMQIFVKTL
TGKTITLEVE PSDTIENVKA KIQDKEGIPP DQQRLIFAGK QLEDGRTLSD YNIQKESTLH
LVLRLRGGMQ IFVKTLTGKT ITLEVEPSDT IENVKAKIQD KEGIPPDQQR LIFAGKQLED
GRTLSDYNIQ KESTLHLVLR LRGGMCENLY FQGHHHHHHH H

*FlexUB6*

MQIFVKTLTG KTITLEVEPS DTIENVKAKI QDKEGIPPDQ QRLIFAGKQL EDGRTLSDYN
IQKESTLHLV LRLRGGSGGS GGSGGSGGMQ IFVKTLTGKT ITLEVEPSDT IENVKAKIQD
KEGIPPDQQR LIFAGKQLED GRTLSDYNIQ KESTLHLVLR LRGGSGGSGG SGGSGGMQIF
VKTLTGKTIT LEVEPSDTIE NVKAKIQDKE GIPPDQQRLI FAGKQLEDGR TLSDYNIQKE
STLHLVLRLR GGSGGSGGSG GSGGMQIFVK TLTGKTITLE VEPSDTIENV KAKIQDKEGI
PPDQQRLIFA GKQLEDGRTL SDYNIQKEST LHLVLRLRGG SGGSGGSGGS GGMQIFVKTL
TGKTITLEVE PSDTIENVKA KIQDKEGIPP DQQRLIFAGK QLEDGRTLSD YNIQKESTLH
LVLRLRGGSG GSGGSGGSGG MQIFVKTLTG KTITLEVEPS DTIENVKAKI QDKEGIPPDQ
QRLIFAGKQL EDGRTLSDYN IQKESTLHLV LRLRGGSGGS GGSGGSGGWC ENLYFQGHHH
HHHHH

*NEMOUBAN6*

MHHHHHHHHE NLYFQGCWGG SMQLEDLKQQ LQQAEEALVA KQEVIDKLKE EAEQHKIVME

TVPVLKAQAD IYKADFQAER QAREKLAEKK ELLQEQLEQL QREYSKLKAS SQESGGSGGS

GGSGGSMQLE DLKQQLQQAE EALVAKQEVI DKLKEEAEQH KIVMETVPVL KAQADIYKAD

FQAERQAREK LAEKKELLQE QLEQLQREYS KLKASSQESG GSGGSGGSGG SMQLEDLKQQ

LQQAEEALVA KQEVIDKLKE EAEQHKIVME TVPVLKAQAD IYKADFQAER QAREKLAEKK

ELLQEQLEQL QREYSKLKAS SQESGGSGGS GGSGGSMQLE DLKQQLQQAE EALVAKQEVI

DKLKEEAEQH KIVMETVPVL KAQADIYKAD FQAERQAREK LAEKKELLQE QLEQLQREYS

KLKASSQESG GSGGSGGSGG SMQLEDLKQQ LQQAEEALVA KQEVIDKLKE EAEQHKIVME

TVPVLKAQAD IYKADFQAER QAREKLAEKK ELLQEQLEQL QREYSKLKAS SQESGGSGGS

GGSGGSMQLE DLKQQLQQAE EALVAKQEVI DKLKEEAEQH KIVMETVPVL KAQADIYKAD

FQAERQAREK LAEKKELLQE QLEQLQREYS KLKASSQES

*UBA6 (from Ubiquilin)*

MQNPEVRFQQ QLEQLSAMGF LNREANLQAL IATGGDINAA IERLLGSGGS GGSGGSGGSQ

NPEVRFQQQL EQLSAMGFLN REANLQALIA TGGDINAAIE RLLGSGGSGG SGGSGGSQNP

EVRFQQQLEQ LSAMGFLNRE ANLQALIATG GDINAAIERL LGSGGSGGSG GSGGSQNPEV

RFQQQLEQLS AMGFLNREAN LQALIATGGD INAAIERLLG SGGSGGSGGS GGSQNPEVRF

QQQLEQLSAM GFLNREANLQ ALIATGGDIN AAIERLLGSG GSGGSGGSGG SQNPEVRFQQ

QLEQLSAMGF LNREANLQAL IATGGDINAA IERLLGSGGS GGSGGSGGSW CENLYFQGHH

HHHHHH

**S1. Supplementary figure 1**

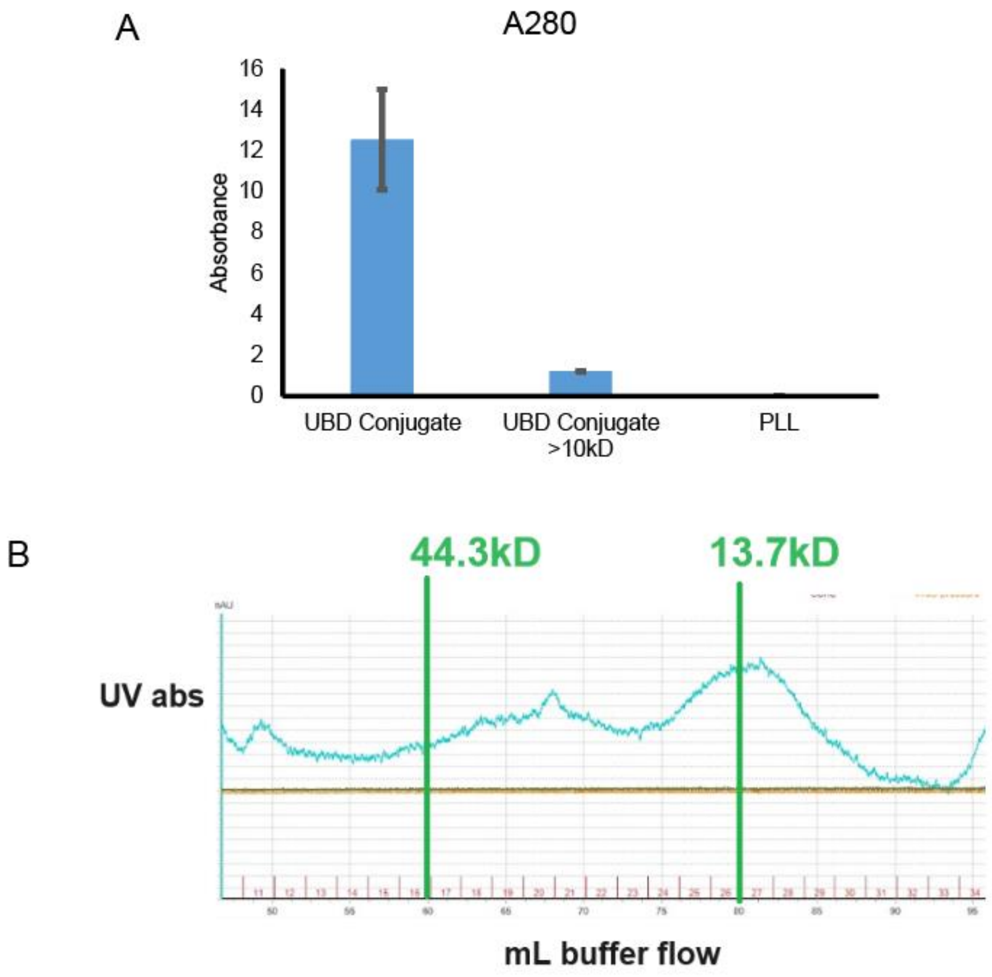


**Figure 1.** Molecular weight characterization of the UBD Conjugate. (A) Nanodrop UV absorption shows that PLL has no UV absorption at all and about 10% of the original UBD peptide end up participating in the crosslinking reaction to make final molecuarweight > 10kD. (B) Superdex75 automated size exclusion analysis of the UBD conjugates shows the size is distributed around 10-40kD range.

### S2. Supplementary figure 2

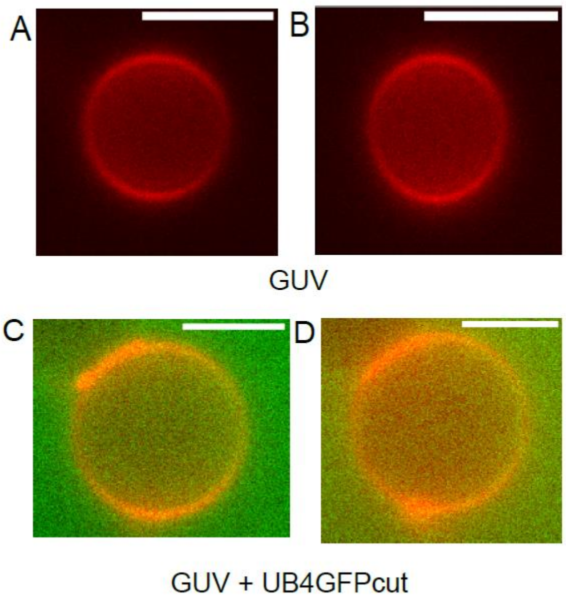


**Figure S2.** Negative control of UB4GFP_cut interaction with GUVs. (A, B) Example images of homogenous GUVs in Texas Red lipid fluorescence (red) channel. (C, D) Example images of unbound UB4-GFPcut cargo, which is a UB4-GFP construct where his-tag was removed by TEV digestion and affinity chromatography. Background fluorescence (green) shows with no colocalization with Texas Red lipid fluorescence (red), representative of unbound protein. The composition of the GUVs used for this control was DOPC 25%, DPPC 39.8%, cholesterol 25%, Ni-DGS 10%, and TR-DHPE 0.2%. It proves that the proteins are not capable of binding to the membranes without the Ni-His tag interaction.

### S3. Supplementary figure 3

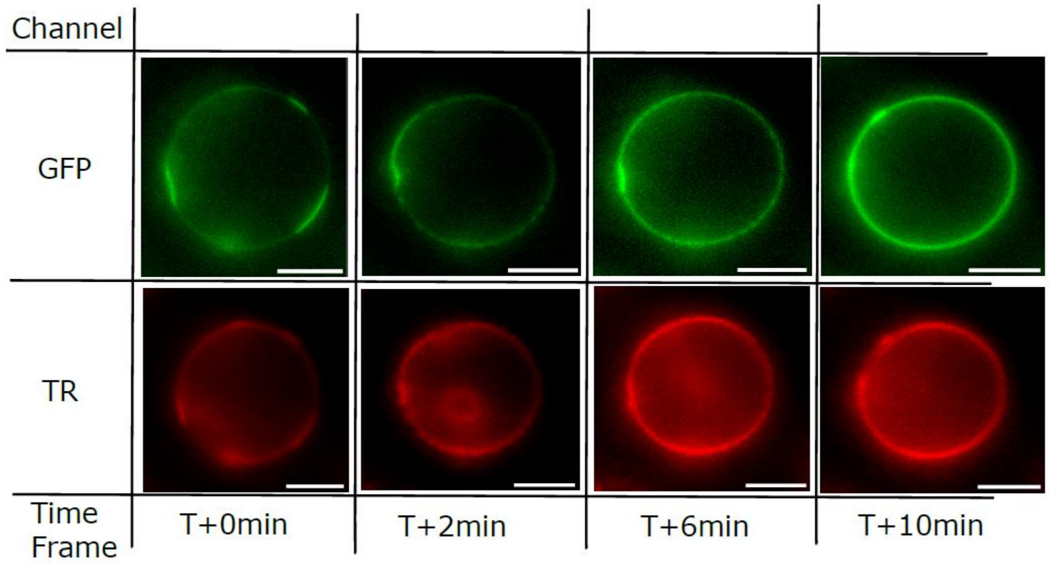


**Figure S3.** Time lapse images show dynamic changes of the phase separation states in Texas Red and GFP channel of a GUV and UB4GFP and UBA6. Time lapse taken at T+0min, 2min, 6min, and 10min. Images show an example kinetic trace reversal of phase separated GUVs becoming homogenous after the addition of the UBA6 at time zero. The composition of the GUVs used for this control was DOPC 25%, DPPC 39.8%, cholesterol 25%, Ni-DGS 10%, and TR-DHPE 0.2%. It proves that the proteins are not capable of binding to the membranes without the Ni-His tag interaction, and sequential addition of UB4-GFP and UBA6 (>10µM) was performed similarly to the main experiment.

### S4. Supplementary figure 4

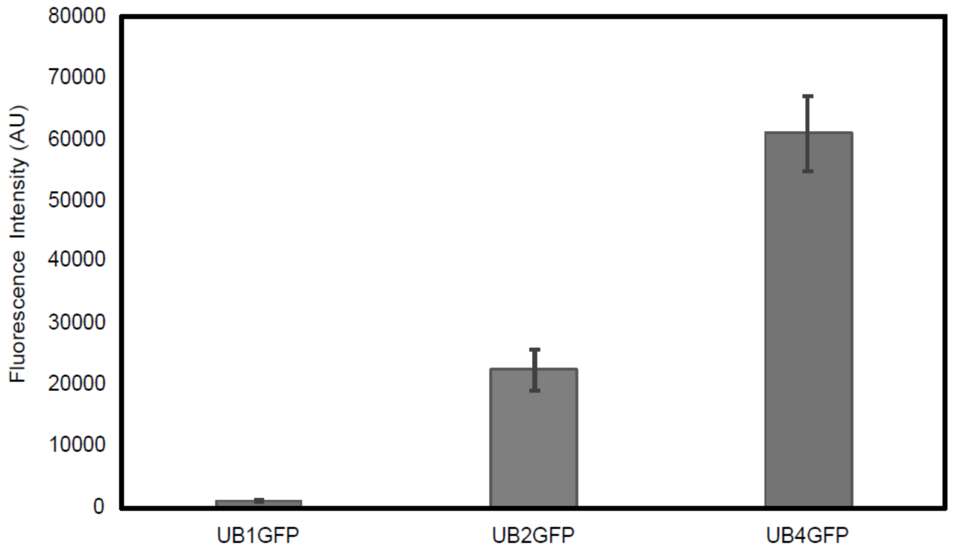


**Figure S4.** Statistical distribution of the maximum fluorescence intensity of droplet condensates formed by UB proteins and UBD Conjugate in solution. Peak fluorescence intensity at the middle of each droplet was quantified for comparison. Error bars represent standard deviations of droplet intensity values of at least 50 unique droplets from 3 z-stack images for each UB cargo protein. It is clear that longer UB chain-based droplets are more enriched with the cargo fluorescence.

### S5. Supplementary figure 5

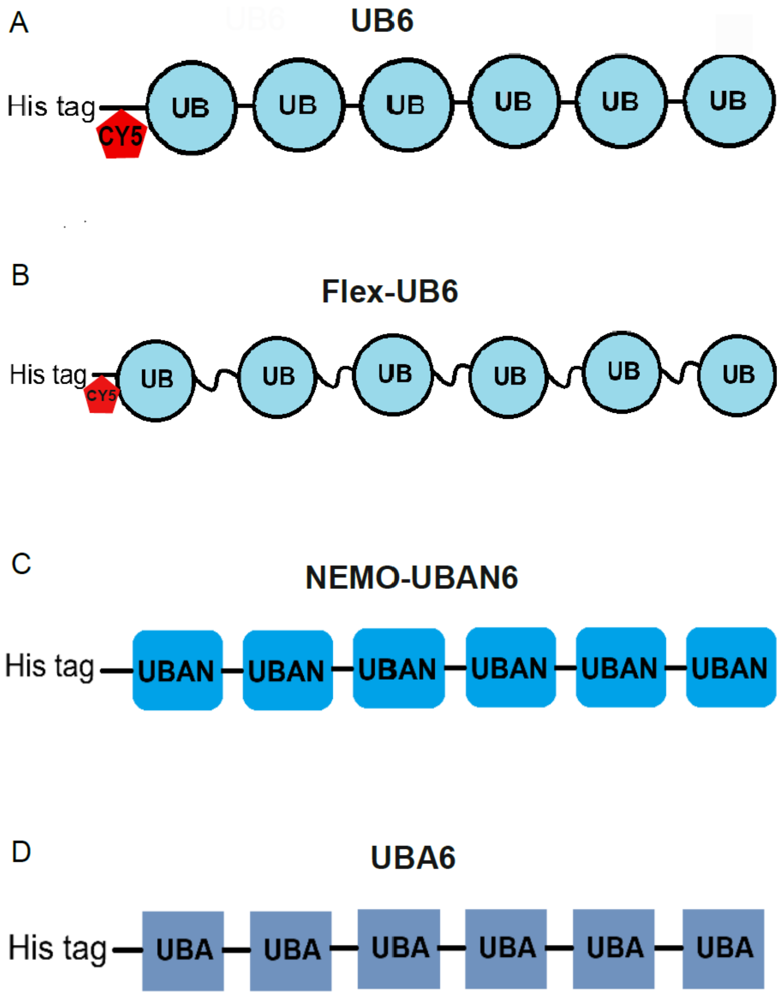


**Figure S5.** Schematics of other modular UB proteins. (A) UB6, polyUB membrane cargos were created by purifying His-tagged repetition of six linear polyUB with a cysteine residue for Cy5 dye attachment. (B) FlexUB6, polyUB membrane cargos were created by purifying His-tagged repetition of six linear polyUB with flexible -GGS- linkers and a cysteine residue for Cy5 dye attachment. (C) NEMOUBA6, Six linear UBAN domains. (D) UBA6, Six linear UBA domains from Ubiquilin.
